# Cooperative effects in DNA-functionalized polymeric nanoparticles

**DOI:** 10.1101/2025.04.06.647366

**Authors:** Paraskevi Gaki, Andrey S. Klymchenko

## Abstract

DNA-functionalized nanoparticles (NPs), called as spherical nucleic acids (SNA), attract strong attention due to their unique properties and numerous applications. In particular, DNA-functionalized dye-loaded polymeric NPs (DNA-NPs), owing to their exceptional fluorescence brightness emerge as powerful nanomaterials for ultrasensitive detection and imaging of nucleic acids. Here, we addressed a fundamental question unexplored for polymeric DNA-NPs: how dense packing of oligonucleotides at the particle surface impacts their capacity to specifically hybridize with complementary sequences. Using Forster Resonance Energy Transfer (FRET) between DNA-NPs and labelled complementary strands, we found that the DNA at the NPs surface exhibit dramatic enhancement in the duplex stability compared to free DNA duplexes (>20 °C). This effect increases at higher densities of DNA on NPs surface, which suggests that DNA cooperativity is responsible for the duplex stability enhancement. For example, 8 nt DNA duplexes were perfectly stable at RT on the surface of DNA-NPs. Furthermore, these DNA-NPs preserve capacity to distinguish mutations, even at the single nucleotide level within 21 nt sequence, when appropriate hybridization temperature is used. The hybridization between DNA-NPs and the complementary sequences proceeds on the min time scale at probe and target concentrations ≥ 10 and ≥100 pM, respectively. Below that, this diffusion-controlled processes becomes too slow, which points on the fundamental limitation in the DNA/RNA sensing assays that require sufficiently high nanoprobe concentration. The present study sheds light on the capacity of DNA-NPs to specifically hybridize with the target sequences and provides insights in the development of nucleic acid sensing assays.

## Introduction

Luminescent nanoparticles (NPs) gained important interest due to multiple unique characteristics, such as high surface to volume ratio, multi-functionality of their surface, capacity to encapsulate large quantities of cargo, etc.^1-2^ Their exceptional brightness provides high signal to noise ratio in biosensing and bioimaging important for diagnostics applications.^3-4^ Luminescent NPs nanoparticles could be split into inorganic, such as quantum dots (QDs),^5^ dye-loaded silica NPs,^6^ metal nanoclusters,^7^ metal-organic framework NPs,^8^ carbon dots,^9^ etc and organic, such as conjugated polymer NPs,^10-12^ aggregation-induced emission (AIE) NPs,^13-15^ dye-loaded polymer^16-17^ and lipid^18^ NPs. Among them dye-loaded polymeric NPs are of particular interest because they provide high stability in biological media, good capacity to encapsulate functional cargoes and rich surface chemistry and biocompatibility.^16,19-20^ Early reports showed that polymeric NPs could encapsulate conventional dyes only at ∼1 wt%, whereas higher loadings lead to aggregation-caused quenching and thus loss in the particle brightness.^19,21-22^ We addressed this problem by proposing bulky hydrophobic counterions as insulators preventing dye aggregation and ensuring efficient loading without dye leakage in biological media.^23-24^ This approach ensured efficient dye emission even at dye loadings reaching >30 wt% of the particle mass, which resulted in exceptional particle brightness^4^ and unique light-harvesting properties,^25^ important for bioimaging^26-27^ and biosensing.^28-30^

A particular class of NPs are those coated with nucleic acids, which Mirkin and co-workers coined as spherical nucleic acids (SNA).^31-32^ These are spherical nanoparticles coated with nucleic acids at high density, which renders them unique properties, such as programmable assembly, high stability against enzymes and high colloidal stability. These properties led to preparation of new type of self-assembled materials as well as led to new technologies for nucleic acid sensing (molecular diagnostics) and gene regulation, precision medicine and immunotherapy.^32-34^ The majority of early examples of SNA were based on metallic (mainly gold) NPs,^31-32^ whereas polymeric NPs functionalized with DNA were reported only in 2018 by Mirkin and co-workers for PLGA.^35^ and by our group for PMMA derivative.^28^ A lot of efforts were focused on preparation of DNA-polymer conjugates,^36^ but the modification of hydrophobic polymers in form of NPs with DNA remained challenging. We reported preparation of DNA-functionalized polymeric NPs, which included special polymer design that ensures high exposure of azide groups of NPs surface.^28^ The latter enabled effective grafting of short DNA at high density, which yielded polymer-based SNA. Due to high brightness and light-harvesting properties they enabled amplified FRET-based detection of DNA fragments^28^ and microRNA,^37^ with sensitivity down to single molecule hybridization,^29^ using robust ratiometric output, compatible with smartphone detection.^38^ More recently, these NPs enabled amplified fluorescence in situ hybridization for RNA imaging in cells,^39^ as well as zeptomole-sensitivity detection of viral RNA in combination with magnetic beads.^30^

The intriguing question in the field of SNA is their cooperativity and multivalence due to presence of numerous repeats of DNA with the same orientation and high local concentration. Multivalency in NPs is a generic phenomenon,^40^ which is known to increase their affinity to corresponding targets by multiple presentation of small molecular ligands,^41-43^ protein ligands,^44^ antibodies,^45^ and it was extensively supported by the theoretical modeling.^46-47^ Regarding SNA, the early studies on SNA based on gold NPs suggested that higher density of oligonucleotides on SNA surface led to higher duplex stability (higher melting points, T_m_) and sharper melting transition curves due to cooperative effects.^48^ Overall, the gain in the T_m_ values was ∼5 °C for gold SNA vs free DNA duplex.^31^ Further studies on cross-linked micellar SNA reported an increase in the T_m_ by 16.5 °C for DNA duplexes between two SNA vs free DNA duplex,^49^ although hybridization of multiple duplexes between the two NPs probably contributed to this significant T_m_ raise. To the best of our knowledge, the multivalency effect has not been well addressed for DNA-functionalized polymeric NPs. In this respect, we were particularly interested to explore the extend of the DNA cooperativity effect in dye-loaded polymeric NPs, the impact of the surface density of the coding sequence, specificity of the hybridization and the sensitivity to mutations. All these questions are essential for further applications of polymeric SNA in nucleic acid detection and imaging applications, where these NPs have already shown a significant promise.^26-30^

In the present work, we addressed a fundamental unexplored question on how high density of oligonucleotides grafted at the surface of polymeric DNA-NPs impacts their capacity to specifically hybridize with complementary sequences and how stable are obtained duplexes. Using FRET approach, we found the remarkable enhancement in the stability of DNA duplexes on the surface of polymeric NPs caused by the DNA cooperativity effect. DNA-NPs can distinguish well a single mutation in the complementary sequences and hybridize at the min time scale when the nanoprobe/target concentrations at high pM level (10-100 pM). These results show strong potential of DNA-NPs as DNA/RNA nanoprobes and provide insights for development of next generation detection assays.

## Materials and methods

### Materials

Rhodamine B octadecyl ester tetrakis-(pentafluorophenyl)borate (R18/F5-TPB) was synthesized by ion ex-change and purified by column chromatography as explained previously.^24^ Polymer PEMA-AspN3-5% was synthesized as described previously.^29^ Sodium phosphate dibasic dihydrade (>99.0%) and sodium phosphate monobasic (>99.0%), both were purchased from Sigma-Aldrich and used to prepare 20 mM phosphate buffers at pH 7.4. Millex^®^-GP Filters (pore size 0.22 μm, diam. 33 mm) and Amicon Centrifugal filters (0.5 mL, 100 kD) were purchased from Sigma-Aldrich. Ultrapure DNase/RNase-Free water (Invitrogen) and was used in all the experiments. Dulbecco’s Phosphate Buffered Saline (PBS) and low binding microcentrifuge tubes were purchased from Dutscher.

### Nucleic Acids Sequences

Lyophilized single strand DNA and RNA sequences were purchased from Biomers GmbH and Eurogentec, dissolved in Ultrapure DNase/RNase-Free water, aliquoted, and stored at −20 °C for further experiments. The oligonucleotide sequences used in this study are shown below.

Sur NPs: 5’ CCC-AGC-CTT-CCA-GCT-CCT-TGA - (DBCO) 3’

A20-DBCO: 5’ AAA-AAA-AAA-AAA-AAA-AAA-AA - (DBCO) 3’

Acceptor-8: 5’ (Atto 665) – TCA-AGG-AG 3’

Acceptor-12: 5’ (Atto 665) – TCA-AGG-AGC-TGG 3’

Acceptor-21: 5’ (Atto 665) – TCA-AGG-AGC-TGG-AAG-GCT-GGG 3’

Acceptor-21_1MUT: 5’ (Atto 665) - TCA-AGG-AGC-TCG-AAG-GCT-GGG 3’

Acceptor-21_3MUT: 5’ (Atto 665) – TCA-AAG-AGC-TCG-AAG-GGT-GGG 3’

Donor-21: 5’ CCC-AGC-CTT-CCA-GCT-CCT-TGA - (Cy3) 3’

### Synthesis of fluorescent nanoparticles

50 μL of the polymer (PEMA-AspN3-5%) solution in acetonitrile (2 mg mL^−1^ containing R18/F5-TPB at 50 wt.% relative to the polymer, meaning 33 wt% to particle mass, was added quickly using a micropipette to 450 μL of 20 mm phosphate buffer (PB), pH 7.4 at 21 °C under shaking (Thermomixer comfort, Eppendorf, 1150 rpm). Then, the residues of acetonitrile were evaporated under reduced pressure.

### Synthesis of DNA-functionalized nanoparticles (DNA-NPs)

Aliquots of DBCO-sequence (20 μM in the final reaction mixture) were added to 100 μL of corresponding nanoparticles. In the case of NPs with different percentage of coding sequence, the total concentration was 20 μM. They were then mixed carefully, microcentrifuged and kept overnight (21 h) at 40 °C in the incubator chamber without shaking protected from light. Afterwards, the mixtures were cooled down to room temperature. These reaction mixtures were purified by centrifugation using centrifuge filters (Amicon, 0.5 mL, 100 kDa) on 1000 rcf at 20 °C for 5 min. The procedure of centrifugation was repeated 4 times to remove the non-reacted oligonucleotides. The obtained DNA-functionalized nanoprobes were diluted to 1 nM concentration and were kept in the dark at 4 °C.

### Characterization of Nanoparticles

Dynamic light scattering (DLS) measurements were performed on a Zetasizer Nano ZSP (Malvern Instruments S.A.). The Zetasizer software provided with standard cumulates and size distribution by volume analysis was used to characterize nanoparticles by DLS. For the data analysis, the following parameters were used: for the solvent (water) – temperature 25 °C, refractive index RI 1.33, and viscosity 0.8872 cP. Nanoparticles were assumed to be all homogenous and spherical in shape. Absorption spectra were recorded on a Cary 5000scan UV–vis spectrophotometer (Varian). Emission spectra were recorded on an FS5 Spectrofluoremeter (Edinburg Instruments) and with FluoroMax-4 Spectrofluorometer (HORIBA Scientific) equipped with 350B Thermoelectric Temperature Controller (Newport) for the measurements requiring strict temperature control. For standard recording of fluorescence spectra, the excitation wavelength was set at 530 nm. The fluorescence spectra were corrected for detector response and lamp fluctuations.

### DNA cooperativity study on the surface of DNA-NPs

For FRET between DNA-modified NPs and different concentrations of acceptor-bearing sequence of 21 nucleotides, the aliquots of Acceptor-21 of varying concentrations were added to 100% coding (100% survivin capture sequence) or 10% coding (10% survivin capture sequence & 90% A20 sequence) NPs (50 wt% of R18/F5-TPB dye with respect to the polymer, meaning 33 wt% of the total mass of the NPs) in Mg Buffer, to a final concentration of 100 pM of NPs in final volume of 300μL, in Low Binding microcentrifuge tubes. The Mg buffer contained 20 mM of Phosphate Buffer along with 12 mM MgCl_2_ and 30 mM NaCl at pH 7.4. The mixtures were incubated at 40°C for 20 minutes protected from light. Afterwards they were cooled down to room temperature. Finally, steady state spectra were recorded.

### Thermal stability of duplexes of DNA-NPs with acceptor-sequences

For thermal stability studies using FRET between DNA-modified NPs and acceptor-bearing sequence, the aliquots of Acceptor-8 or Acceptor-12 at 10 nM final concentration were added to 100% coding (100% survivin capture sequence) or 10% coding (10% survivin capture sequence & 90% A20 sequence) NPs (50 wt% of R18/F5-TPB dye with respect to the polymer, meaning 33 wt% of the total mass of the NPs) in Mg Buffer, to a final concentration of 100 pM of NPs in final volume of 600μL, in Low Binding microcentrifuge tubes. The mixtures were incubated at 40°C for 20 minutes protected from light. Afterwards they were cooled down to room temperature. For the mixtures, a fluorescent spectrum was recorded at the starting point of 20°C and every 2°C until 60°C (for the samples with acceptor-8) or until 80°C (for the samples with acceptor-12). The temperature was increased gradually using a Peltier-based temperature control, incorporated in the fluorometer.

### Thermal stability of individual DNA duplexes

For thermal stability studies using FRET between a single dye donor attached to the same DNA sequence that is present on the NPs (Donor-21) and acceptor-bearing sequence, the aliquots of Acceptor-8 or Acceptor-12 at 10 nM final concentration were added to Donor-21 at 3.3 nM final concentration in Mg Buffer, to a final volume of 600μL, in Low Binding microcentrifuge tubes. The mixture containing Acceptor-8 was incubated at 4°C for 1 hour, whereas the mixture containing Acceptor-12 was incubated at 40°C for 20 minutes, both protected from light. The latter was cooled down to room temperature before starting measurements. For the mixture with Acceptor-8, a temperature range from 6°C to 40°C was selected, and for the mixture with Acceptor-12, a temperature range from 20°C to 60°C was selected. A fluorescent spectrum was recorded at the lowest temperature and every 2°C until the highest. The temperature was increased gradually using a Peltier-based temperature control, incorporated in the fluorometer. The melting temperature for the 8mer and 12mer was estimated using online calculator (http://insilico.ehu.es/, which took into account DNA concentration (10 nM) and composition of Mg buffer.

### Sensitivity to mutations of DNA-NPs

To study the sensitivity to mutations of the NPs, FRET between DNA-modified NPs and acceptor-bearing sequence with zero, one and three point mutations was used. Aliquots of Acceptor-21, Acceptor-21_MUT1 and Acceptor-21_MUT3 at 10 nM final concentration were added to 100% coding (100% survivin capture sequence) or 10% coding (10% survivin capture sequence & 90% A20 sequence) NPs (50 wt% of R18/F5-TPB dye with respect to the polymer, meaning 33 wt% of the total mass of the NPs) in Mg Buffer, to a final concentration of 100 pM of NPs in final volume of 600μL, in Low Binding microcentrifuge tubes. The mixtures were incubated at 40°C for 20 minutes protected from light. Afterwards they were cooled down to room temperature. For the mixtures, a fluorescent spectrum was recorded at the starting point of 20°C and every 2°C until 70°C. The temperature was increased gradually using a Peltier-based temperature control, incorporated in the fluorometer.

### Kinetics of hybridization of DNA-NPs with acceptor-sequences

Aliquots of Acceptor-21 were added to 100% coding (100% survivin capture sequence) NPs (50% wt% of R18/F5-TPB dye with respect to the polymer) in Mg Buffer, to a final volume of 600μL, in the plastic cuvettes that are used to record the spectra, as to not waste time. The components were used in 1 NP: 10 Acceptor ratio, at different concentrations, starting from 1 pM NPs mixed with 10 pM Acceptor-21. Fluorescent spectra were recorded for 10 minutes with the rate of one spectrum per minute. Before proceeding to any analysis, all spectra were smoothed by using Adjacent-Averaging with 5 Points of Window.

### FRET ratio calculation

FRET ratio was calculated using the following equation:

FRET ratio = I_A_ / (I_A_ + I_D_) where I_A_ is the fluorescence intensity of the acceptor and I_D_ is the fluorescence intensity of the donor (NPs or single dye donor).

## Results and Discussion

### DNA-NPs and their hybridization studied by FRET

The designed DNA-NPs are comprised of a dye-loaded polymeric core and a single-stranded DNA shell. The core is based on polyethyl methacrylate-co-methacrylic acid bearing aspartic acid with azide (PEMA-AspN3, 5% methacrylic acid)^29^ and is loaded at 33 wt% (to particle mass) with an ion pair of hydrophobic rhodamine (R18) with the bulky hydrophobic counterion tetrakis(pentafluorophenyl)borate (F5-TPB) to avoid aggregation-caused quenching (Figure 1). The dye molecules of the core act as the energy donor in a Förster Resonance Energy Transfer (FRET) pair and can transmit energy to an acceptor dye on the surface.^28-29^ To obtain DNA-NPs, the NP core bearing azide groups is functionalized with a single-stranded capture oligonucleotide bearing DBCO by stain promoted cycloaddition reaction (Figure 1).^28-29^ DNA fragment of survivin, which is an inhibitor of apoptosis protein that is widely expressed in malignant cells and an important cancer marker, was selected as a target complementary to DNA capture sequence of DNA-NPs. In order to study how the composition of the DNA shell of the NPs affects the hybridization with the complementary target sequence, we exploited FRET from NP core to acceptor-modified target hybridized at the NPs surface to the capture oligonucleotide (Figure 1).

**Figure 1.**
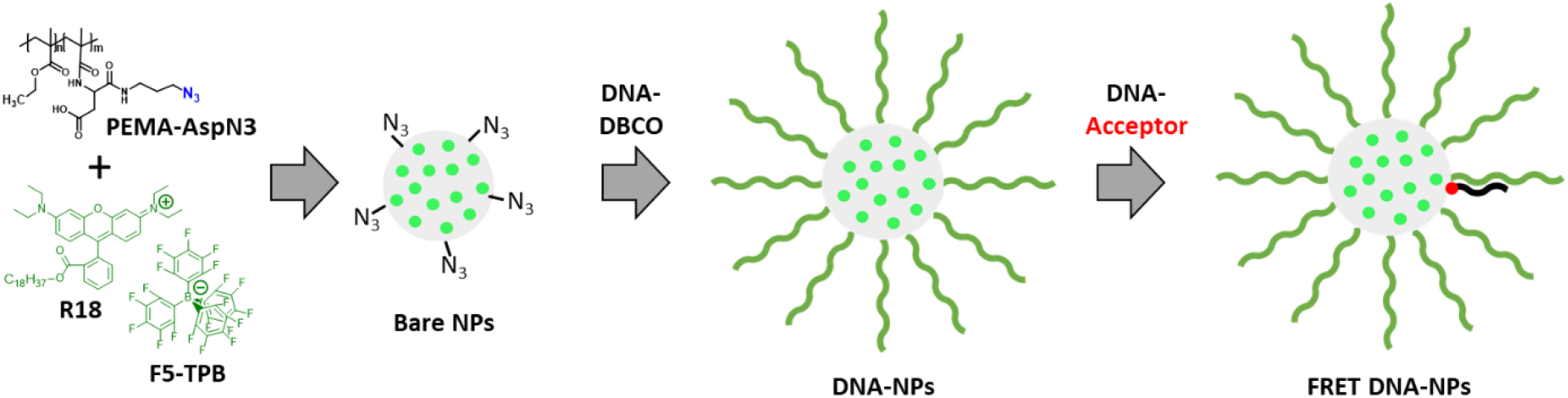
Chemical structures of PEMA-AspN3 polymer and dye R18/F5-TPB and their nanoprecipitation into bare dye-loaded polymeric NPs with functional azide groups and then functionalization of the latter into DNA-NPs and hybridization of the complementary stands with FRET acceptor.

The core of the nanoparticle was prepared by nanoprecipitation of PEMA-AspN3 and R18/F5-TPB to yield NPs of 33 nm size and good polydispersity according to DLS (Table S1). TEM confirmed spherical shape and small size (23 ± 4 nm) of the obtained NPs (Figure S1). They showed a single absorption and emission spectra typical for R18 dye (Figure S2). Then, the obtained NPs were functionalized with different fractions of coding sequence. As coding sequence on the NPs, the single stranded DNA complementary to a part of the survivin mRNA, an important cancer marker, was selected. As non-coding oligonucleotide, a single stranded A20 was used. The NPs were prepared with 100% and 10% of coding sequence and 0% and 90% of A20 sequence, respectively. After functionalization with oligonucleotides, their hydrodynamic diameter increased by 12-13 nm (Table S1), which corresponded to the presence of the additional DNA shell. TEM showed similar spherical shape (Figure 2 and S3) and slightly increased diameter: 28 ± 5 and 32 ± 6 and for 100 and 10% coding DNA-NPs, respectively (Table S1). Their absorption and emission spectra remained unchanged after functionalization (Figures S4-S5). The fluorescence quantum yield of obtained DNA-NPs (10% coding) was 53%, which was in line with earlier works on PEMA-AspN3 NPs.^29^

**Figure 2.**
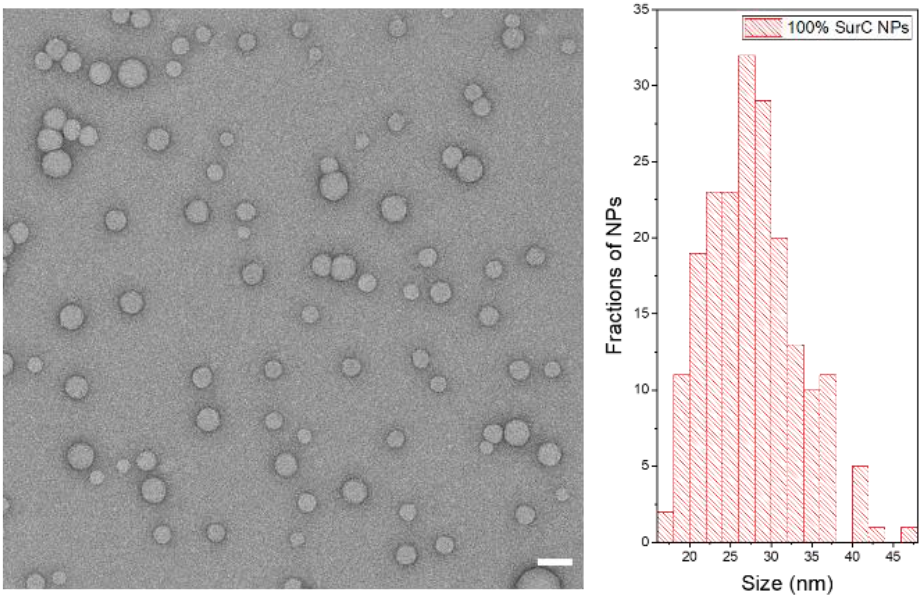
TEM image (left) and size distribution diagram (right) PEMA-AspN3 NPs functionalized at 100% with survivin capture coding sequence and loaded with 33 wt% of R18/F5-TPB dye (with respect to total NP mass). Scale bar: 50 μm.

Increasing concentration of target (complementary) DNA sequence labeled with an acceptor dye (Atto-665) was added to each type of NPs and the fluorescence spectra were recorded (Figure 3A). For all studied NPs, the increase in the DNA-acceptor concentration led to higher signal of the long-wavelength band corresponding to the FRET acceptor. Using the fluorescence spectra, the FRET ratios were calculated for all conditions, using the formula I_A_ / (I_A_ + I_D_), where I_A_ and I_A_ are the peak intensities of the acceptor and donor, respectively, in the fluorescence spectrum. FRET ratio gives a semi-quantitative information on the FRET efficiency and consequently the efficacy of hybridization between the capture DNA on the NPs and the target. For both studied NPs, a steep growth of the FRET ratio was observed upon increase in the DNA-acceptor concentration. At higher DNA-acceptor concentrations, the curves reached a plateau, but the saturation values of the FRET ratio were higher for the NPs with higher fraction of the coding sequence. For 10 and 100% coding DNA-NPs, the observed saturations occurred at 2 and 5 nM DNA-acceptor, respectively. This saturation suggested that the DNA-NPs reached their maximum capacity to hybridize DNA-acceptor. Considering that the NPs concentration was 100 pM, and the number of capture DNA per NP is 100-200, we expect the maximal hybridization capacity of 10 and 100% coding DNA-NPs should be achieved at 1-2 and 10-20 nM of DNA-acceptor, respectively. In case of 10% coding DNA-NPs, this estimation matches perfectly the experimental data, suggesting that in this case probably all capture sequences were able to hybridize with the complementary DNA-acceptor. Even though for 100% coding DNA-NPs the saturation was reached at higher concentrations, it was still below the expected value (25-50% of the expected value), suggesting that only 25-50% of capture sequences were hybridized in this case, whereas further hybridization was probably blocked because of steric hindrance produced by increased density produced by newly formed DNA duplexes. The above show that the hybridization capacity depends on the density of the coding nucleic acids: larger fraction of coding sequence helps to hybridize larger amount of the targets, but the hybridization is probably limited by the steric hindrances at the surface, due to high nucleic acid density.

**Figure 3.**
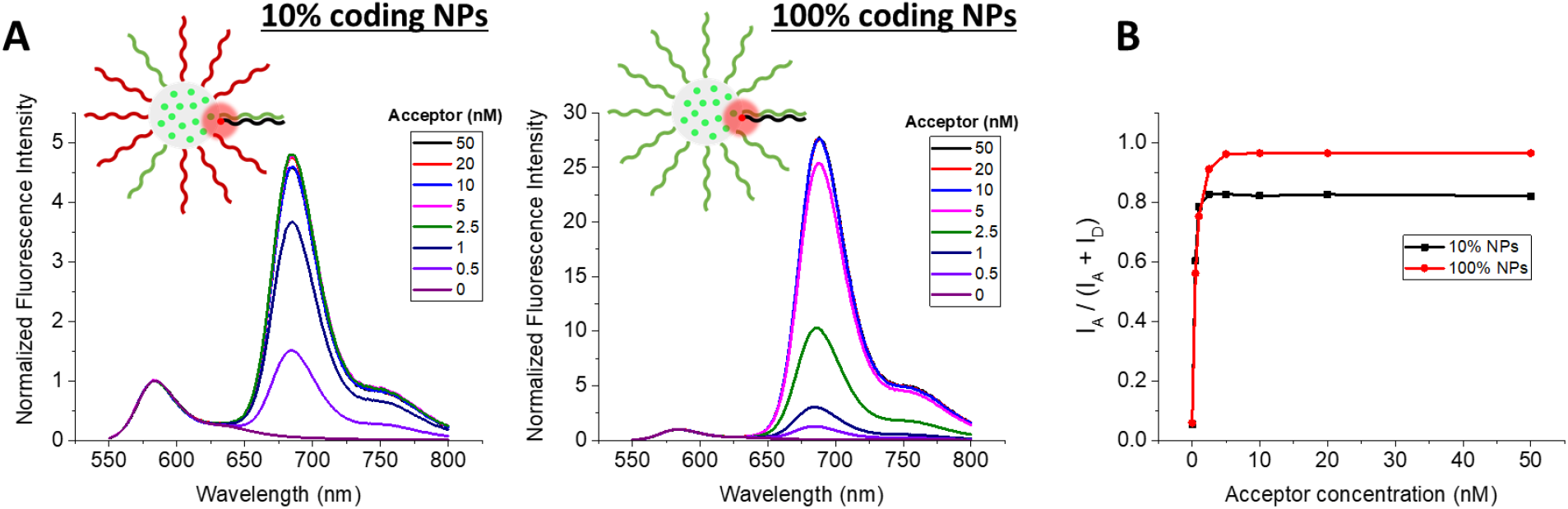
DNA cooperativity effect for DNA-functionalized nanoparticles with varied surface density of coding sequences. A) Fluorescence spectra of 100% coding (100% survivin capture sequence) and 10% coding (10% survivin capture sequence, 90% A20 sequence) NPs (50% wt% of R18/F5-TPB dye with respect to the polymer, meaning 33 wt% of the total mass of the NPs), hybridized with varying concentrations of an acceptor dye bearing DNA sequence of 21 nucleotides, representing part of the survivin target, thus fully complementary to the capture sequence on the NP. For this experiment 100 pM of NPs were hybridized with 0.5 – 50 nM of acceptor-sequence. Negative control: NPs without addition of acceptor-sequence. Mg buffer was systematically used. Spectra measured at 530 nm excitation. B) FRET ratio for the 100% coding and 10% coding NPs, measured by varying the concentration of the acceptor-sequence.

### Thermal stability of the duplexes in DNA-NPs

The high density of the DNA on the NPs surface is expected to lead to cooperative phenomena, which could increase stability of hybridization compared to that of the free single stranded sequences. For this reason, we studied the thermal stability of the duplexes on the surface of the DNA-NPs in comparison to the free DNA duplexes. Two FRET systems were used: (i) DNA-NPs acting as the energy donor and dye-bearing target sequence as the acceptor, and (ii) the DNA capture sequence, as for the DNA-NPs, bearing donor dye and the same target DNA-acceptor. Thus, the DNA sequences used are the same in these two cases and the differences in the hybridization can be directly attributed to the effect of confinement of DNA on the NPs surface.

DNA-NPs were prepared with 100% and 10% coding sequences on their surface as mentioned above. They were hybridized with acceptor dye-labeled target sequences of different lengths: 8 and 12 nt. These short target sequences are complementary to the end attached to the NP, where the acceptor dye is placed next to the donor dye-loaded core of NPs. In the control without NPs, the single stranded DNA bearing the donor dye (Atto 665) was hybridized with acceptor dye-bearing target sequence of 8 and 12 nt. The above-mentioned hybridization was performed by an annealing procedure, where the sample was subjected to a heating step, which disrupts potential secondary structures by breaking all hydrogen bonds, followed by a cooling step, which facilitates the formation of new hydrogen bonds between complementary oligonucleotides.

In order to evaluate the thermal stability of the duplexes we started by calculating the melting temperature for free DNA duplexes (Tm). For the 8mer and 12mer at 10 nM concentration in Mg buffer used, the calculated Tm was 17 and 49°C, respectively. It is important to keep in mind that these are approximations and that the real melting temperature depends on many factors, including but not limited to the buffer composition.

For each duplex sample, a fluorescence spectrum was recorded at the starting point of 20°C and every 2°C until 60-80 °C (Figures 4 and S6-S10). The temperature was increased gradually using a Peltier-based temperature control, incorporated in the fluorometer. The resulting fluorescence spectra were used to calculate the FRET ratios, obtained from the fluorescence spectra. Subsequently, the FRET ratios were plotted against the temperatures to obtain the melting curves. At the starting point for all samples high value of the FRET ratio was observed, indicating that the duplexes were formed. For the free DNA duplex with 8-nt acceptor, the starting temperature was set at 6°C, as the melting temperature was low and we intended to see the transition. The melting curves of free DNA duplexes with 8- and 12-nt acceptors showed the classical melting profiles (Figure 4B). At lower temperatures the FRET ratio remained high, because the sequences were in double stranded form. When arriving close to their expected melting temperature, a decrease in the FRET ratio was observed, signifying the duplex dissociation.

**Figure 4.**
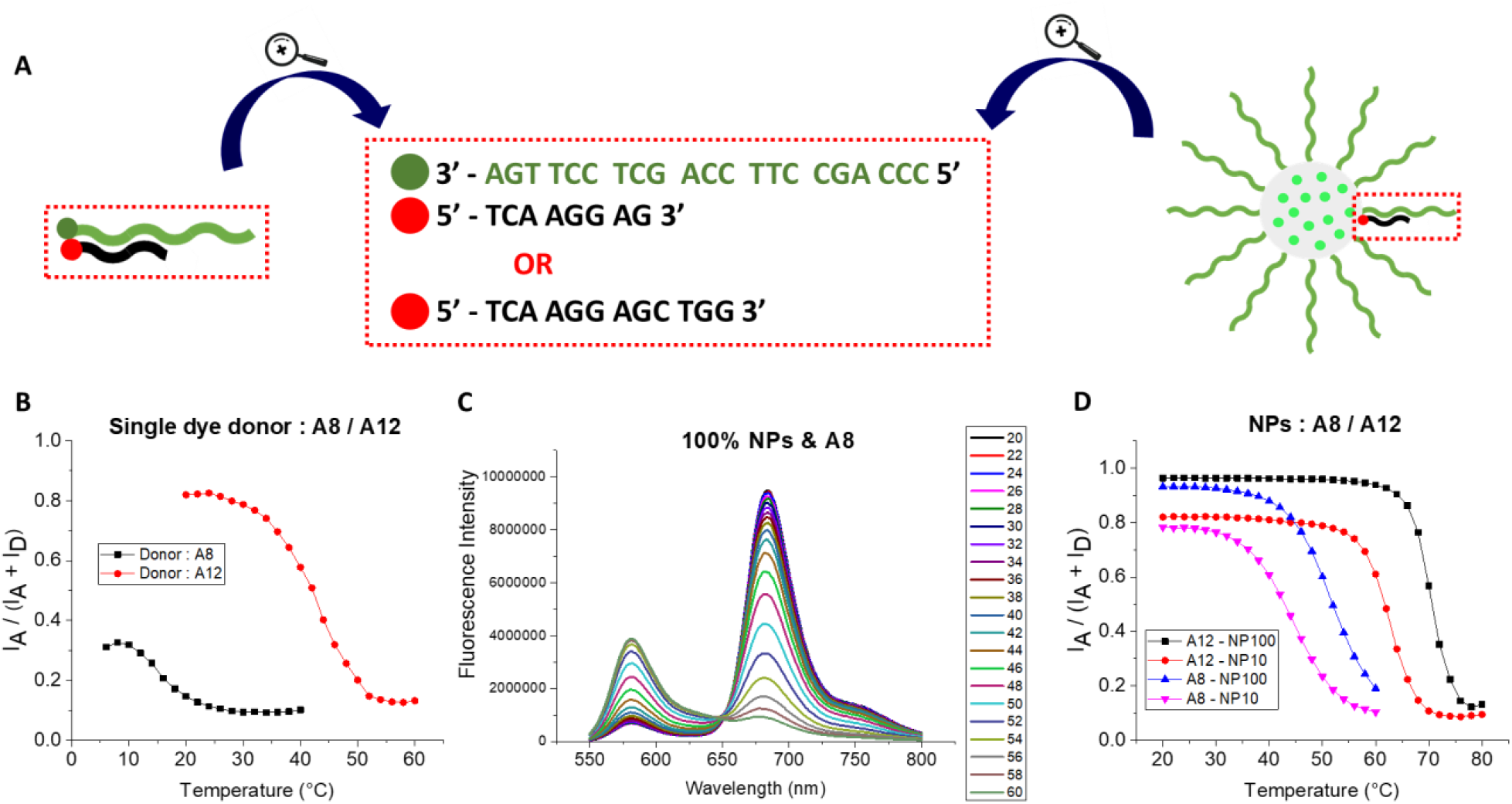
Thermal stability of the duplexes. A) Schematic representation of the single stranded donor and acceptor-sequence pairs and the NP and acceptor-sequence pairs. The oligonucleotide sequences attached to the donor and to the acceptor are presented. B) FRET ratios for increasing temperatures for the pairs of single dye donor: acceptor-8 (in black) and acceptor-12 (in red). For this experiment 3.3 nM of donor-sequence were hybridized with 10 nM of acceptor-sequence. C) Fluorescence spectra for 100% coding NPs hybridized with acceptor-8 and subjected to increasing temperatures from 20°C to 60°C. D) FRET ratios for 10% and 100% coding NPs, hybridized with acceptor-8 and acceptor-12. For this experiment 100 pM of NPs were hybridized with 10 nM of acceptor-sequence. The temperature was increased from 20°C to 60°C (for 10% coding NPs) or to 80°C (for 100% coding NPs). Mg buffer was systematically used. Spectra measured at 530 nm excitation.

In the case of the FRET systems with the DNA-NPs, fluorescence spectra showed strong temperature dependence: initially high signal of the FRET acceptor dropped at higher temperatures, whereas the donor emission increased, indicating the loss of FRET due to disruption of the DNA duplexes (Figure 4C). In the obtained melting curves for 10% and 100% coding DNA-NPs with 8- and 12-nt acceptor, high values of the FRET ratio were observed at lower temperatures (Figure 4D), suggesting the presence of DNA duplexes. Increase in the temperature resulted in a sigmoid response, with sharp decrease in the FRET ratio corresponding to the DNA duplex melting. The melting temperature range was characteristic for each sample, and increased in the following order: 10% DNA-NPs with 8-nt acceptor < 100% DNA-NPs with 8-nt acceptor < 10% DNA-NPs with 12-nt acceptor < 100% DNA-NPs with 12-nt acceptor. Thus, as expected, longer sequences melted at higher temperature. More importantly, increase in the density of coding sequences at the NPs surface increased the melting point (Figure 4D). Another important finding is that melting curves obtained for DNA-NPs were drastically shifted towards higher temperatures compared to free DNA duplexes (Figure 4D). For 8-nt acceptor, 10 and 100% DNA-BPs showed respectively 18 and 26 °C shift of melting curve to higher values compared to corresponding free duplex. For 12-nt acceptor, the melting curve was shifted ∼20 and ∼ 28 °C for 10 and 100% coding NPs, respectively. These results show strong DNA cooperativity effect, where close proximity of coding DNA on NPs surface boosts the duplex stability. The obtained value of Tm enhancement is significantly higher than that reported for classical example of gold SNA, where only enhancement of ∼5 °C was reported for SNA vs free DNA duplex.^48^ The effect is even stronger compared to SNA (to cross-liked micelles) connected by DNA duplexes (16.5 °C).^49^ Moreover, the observed enhancement in the duplex stability with the increase in the density of the coding sequence (by 8 °C from 10 to 100% of coding sequence density) is in line with the earlier report on gold SNA, where Tm increased by 3.1 °C on the increase of coding DNA density from 33 to 100%.^48^ Thus, SNA based on polymeric NPs exhibit remarkably high DNA cooperativity effect, surpassing classical examples of gold SNA, which enables formation of stable duplexes even for relatively short DNA sequences down to 8 nt, which are now stable at RT.

### Sensitivity to mutations

In order to be useful for RNA/DNA sensing in molecular diagnostics, our DNA-NPs should present high level of sequence specificity, ideally with capacity to detect a single mutation in the target sequence. To understand the effect of mutations, the DNA-NPs were investigated in the same FRET-based format. We designed sequences with 1 and 3 point mutations, based on an acceptor dye-labeled target sequence of 21 nt, fully complementary to the capture sequence on the DNA-NPs. The single mutation was introduced at the middle of the sequence (A21_1MUT), whereas the 3 mutations (A21_3MUT) were spread across the sequence keeping approximately the same length in the formed short complementary fragments. This should in theory split the long sequence in shorter ones and decrease the melting temperature.

DNA-NPs with 100% and 10% coding sequences were hybridized with three types of acceptor sequence with different number of mutations. The DNA-NPs were annealed with their complementary sequences labelled with the FRET acceptor, as explained above to achieve complete hybridization. For all studied pairs of DNA-acceptor with DNA-NPs, a fluorescence spectrum was recorded at the starting point of 20°C and then every 2°C until 70°C (Figures S11-S16). At 20°C, the DNA-NPs with the same density of coding sequences presented almost the same high values of the FRET ratio, indicating the presence of stable duplexes (Figure 5). For the 10% coding DNA-NPs, the duplex with the A21_3MUT started melting first, shortly after 40°C, followed by the duplex with the A21_1MUT ∼55°C (Figure 5C). The duplex with the fully complementary sequence showed high stability in the studied temperature range with only the onset of melting at the highest studied temperature (70 °C). When it came to the 100% coding DNA-NPs, the duplex with the A21_3MUT started melting at 50°C, whereas the A21_1MUT duplex started melting after 60°C (Figure 5C). The A21 duplex was stable at the studied temperature range, showing no noticeable changes. Undoubtedly, introducing point mutations decreased drastically the melting temperature for our DNA-NPs, which shows the good sensitivity to mutations.

**Figure 5.**
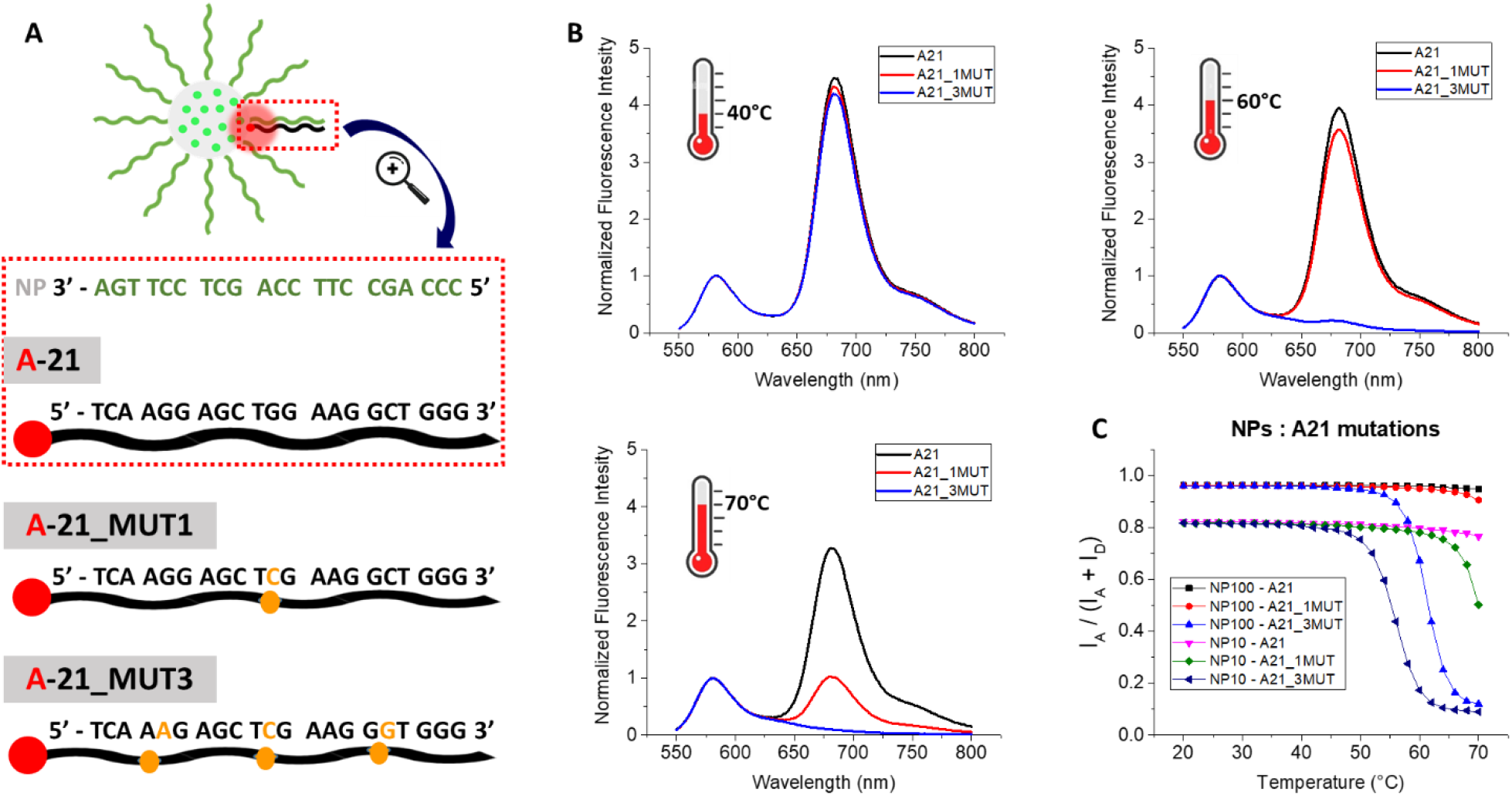
Sensitivity to mutations. A) Schematic representation of the NP hybridized with acceptor-sequences each containing zero (A-21), one (A-21_MUT1) or three (A-21_MUT3) point mutations as indicated with the yellow nucleotides. B) Normalized fluorescence spectra for 10% coding NPs hybridized with acceptor sequences with zero (in black), one (in red) or three (in blue) point mutations at three different temperatures: 40°C, 60°C and 70°C. C) FRET ratios for 10% and 100% coding NPs, hybridized with acceptor-sequences with zero, one or three point mutations. For these experiment 100 pM of NPs were hybridized with 10 nM of acceptor-sequence. The temperature was increased from 20°C to 70°C. Mg buffer was systematically used. Spectra measured at 530 nm excitation.

From the above we can see that both 10 and 100% coding NPs have the capacity to spot mutations, but at a different extent. For the 10% coding NPs, the melting temperatures were systematically lower than for the 100% coding ones, allowing for a more sensitive detection at the normal operational temperature range of the NPs. Indeed, at 60°C, 10% DNA-NPs could clearly distinguish three mutations, whereas at 70°C even a single point mutation was clearly spotted. For the 100% coding NPs, in the working temperature range it is still possible to distinguish the sample with the three mutations, but for the one mutation it cannot be done because of too high melting point.

We come to the conclusion that despite strong DNA cooperativity effects, our DNA-NPs are capable to distinguish mutations, even a single one in a 21nt long oligonucleotide. As a certain temperature range has to be respected because of the nature of the nanoprobes, the 10% coding NPs are a good fit to be used for the detection of sequences with mutations. The DNA-NPs present improved sensitivity to mutations at higher temperatures, but the exact temperature that has to be applied to distinguish one or more mutations has to be defined for each target, as the Tm depends on the sequence.

### Kinetics of hybridization and intrinsic sensitivity of the system

So far, we were focused on the stability of the obtained duplexes of DNA-NPs, which is essentially a thermodynamic parameter. On the other hand, kinetics of the process is also very important, especially in the applications of DNA/RNA sensing, where the target concentration can vary. Therefore, we investigated how fast the DNA-NPs capture their target sequence at room temperature (∼22°C) and what is the lowest concentration of target DNA that can be robustly detected.

In this study, DNA-NPs were mixed with the DNA-Acceptor at different concentrations and the kinetics of the hybridization was monitored as a change in the FRET signal. Based on the results above, 100% coding DNA-NPs and 21-nt DNA-acceptor (A21) were selected, to achieve the most efficient hybridization and the maximal duplex stability. The ratio of DNA-NPs : A21 was kept at 1 : 10, while the concentration of components was decreased gradually down to 1 pM of DNA-NPs and 10 pM of A21. After both components were mixed, the fluorescent spectra of each mixture were recorded immediately and every 1 min until 10 min.

Due to the fast acquisition time required to capture each time point, the spectra were quite noisy for the lowest concentration (Figure 6A). Therefore, the spectra were smoothed in order to calculate properly the FRET ratio, which was then plotted against the time for each sample (Figure 6B). For all studied mixtures, the FRET ratio increased over time, showing that DNA-acceptor hybridized on the DNA-NPs surface, leading to the rise of the FRET signal (Figure 6B). The kinetics of FRET response (i.e. DNA hybridization) showed clear concentration dependence. The rise of the FRET signal was the steepest for the most concentrated sample (37.5 pM of DNA-NPs) with reaction completing within 5 min and the slowest for the most diluted ones (at 1 pM of DNA-NPs). It was observed that the initial speed of hybridization was higher and it decreased over time, which was epically well seen for higher concentrations. Therefore, the kinetics curve could be fitted with a polynomial function (Table S2). It is typical for the bimolecular processes where both components are consumed over time, so that the reaction rate decays non-linearly. Several conclusions can be drawn from these measurements. First, we achieved direct detection of DNA hybridization at concentration down to 10 pM of the DNA target, which is far lower than that possible for typical fluorescence-based molecular hybridization assay (typically in nM range). These became possible due the fluorescence signal amplification through the light-harvesting from our DNA-NPs: the hybridization of a single acceptor leads to >100-fold amplification because of efficient FRET from hundreds of encapsulated dyes to a single acceptor.^25,29^ Measurements at these low concentrations revealed a key feature of the direct hybridization assay: the hybridization between NPs and the complementary sequence (target) is fast (time scale of minutes) and efficient at concentrations of the target and nanoprobe ≥100 and ≥10 pM, respectively. Below these concentrations, the reaction becomes slow, being controlled by the diffusion of the two components. This defines the limit of detection (LoD) of the direct hybridization assay when the kinetics is diffusion controlled. This information will be useful when setting the parameters for a final test as it gives important insight for the lowest concentration of NPs that need to be used and for the minimum time required to complete the hybridization. The obtained results shed light on our previous results, where our FRET assays based on similar DNA-NPs at 10 pM concentration reached LoD of 1-10 pM for the incubation time in the range 1-6 h.^28-29,37^ It shows that when further improvement of sensitivity is needed, the concentration of one of the two hybridization components should be higher than the mentioned limits. As an example, it was recently realized in our direct hybridization assay that reached fM sensitivity to the DNA/RNA targets, where DNA-NPs were used at 200 pM concenrations.^30^

**Figure 6.**
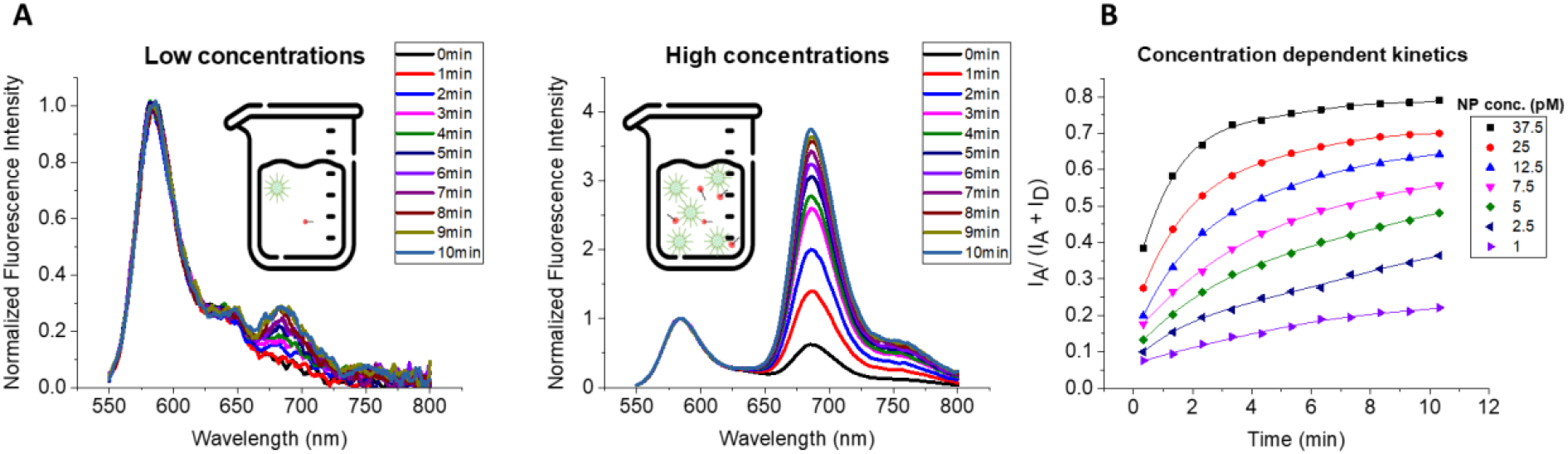
Kinetics of hybridization. A) Normalized (and smoothed) fluorescence spectra for 100% coding NPs mixed with acceptor-sequence with 21 nucleotides at a ratio of 1 NP: 10 acceptor-21 in low and high concentrations. The spectra were recorded right after mixing and for a duration of 10 minutes, with the rate of one spectrum per minute. Schematic representation of the NPs and acceptor-sequences in the solution at different concentrations. C) FRET ratios for varying concentrations of 100% coding NPs: acceptor-21 during 10 minutes. The curves connecting the data point are the polynomial fits. For these experiment the ratio of 1 NP: 10 acceptor-21 was respected throughout, starting with 1 pM of NPs hybridized with 10 pM of acceptor-21 and increasing the concentrations. Mg buffer was systematically used. Spectra measured at 530 nm excitation.

## Conclusions

DNA-functionalized polymeric NPs emerged as powerful nanomaterials within the class of SNA, which, owing to their high fluorescence brightness, enable ultrasensitive detection and imaging of nucleic acids in solutions and cells. Here, we addressed a fundamental unexplored question on polymeric DNA-NPs: how oligonucleotides grafted at the surface of polymeric NPs at high density impact its capacity to specifically hybridize with complementary sequences and how stable are the obtained duplexes. To address it, we employed FRET between DNA-NPs as donors and labelled complementary strands of varied length as acceptor. It was found that the DNA at the NPs surface exhibit dramatic enhancement in the duplex stability compared to free DNA duplexes (>20 °C). This effect increases with increase in the density of DNA on NPs surface, which suggests that a DNA cooperativity effect is responsible for the duplex stability enhancement. As a result, the DNA-NPs were able to capture at RT even short DNA duplexes down to 8 nt, unstable in these conditions. The observed cooperativity effect has been previously shown for gold NPs,^31,48^ although the presently observed temperature enhancement is higher. The phenomenon of DNA cooperativity stems from at least two factors. First, it is high local concentration of DNA grated at high density at the NP surface. This is in line with the fact that the measured melting is lower for 10% coding NPs vs 100% coding NPs. Second, DNA stands are aligned nearly at the NPs surface, which ensures favorable conditions for dehybridized complementarity stand to re-hybridize with the neighbor capture stand. Furthermore, it was shown that DNA-NPs are capable to distinguish mutations, even at the level of the single nucleotide within 21 nt sequence, when appropriate hybridization temperature is used. Previously, Mirkin and co-workers showed that DNA cooperativity makes the melting curves sharper compared to individual DNA strands.^31,48^ This means that at the temperature close to the melting point, DNA-NPs should be even more sensitive to DNA mutations than the individual DNA strands. The hybridization between DNA-NPs and the complementary sequences is fast (min time scale) at probe and target concentrations ≥ 10 and ≥100 pM, respectively. Below that, the reaction takes much longer time, which points on the importance of the use of sufficiently concentrated nanoprobe in the development of fast and sensitive assay for DNA/RNA target detection. These observations explain previous results on FRET-based nanosensors, where the observed limited sensitivity at the low-pM range is linked to slow kinetics of DNA hybridization at too low concentrations of the DNA nanoprobe and nucleic acid target.^28-29,37^ Overall, this work shows that polymeric DNA-NPs present enhanced capacity to hybridize with complementary stands compared to individual DNA and at the same time distinguish well a single mutation when the temperature is respected. Moreover, it provides insights on how to design DNA-NPs in order to obtain better nucleic acid sensing assays.

## Supporting information

Supplemental material for publication

## Acknowledgements

This work was supported by iLab grant MirSens by BPI & Ministère de l’enseignement supérieur et de la recherche, ERC Proof of Concept Grant AmpliFISH 899928, Fondation Force Grant and Région Grand Est collaboration grant VirSens between BrightSens Diagnostics and University of Strasbourg. For Electron microscopy, this work used the Integrated Structural Biology platform of the Strasbourg Instruct-ERIC center IGBMC-CBI supported by FRISBI (ANR-10-INBS-0005).

## Conflicts of interests

ASK is co-founder of BrightSens Diagnostics SAS and PG are currently employed by this company. ASK and PG deposited patent applications related to the described technology.

