## Supplemental material for publication for "Cooperative effects in DNA-functionalized polymeric nanoparticles"

**Table S1.** Hydrodynamic diameter and polydispersity (PdI) by DLS and the diameter by TEM of fluorescent NPs used in this work.

| **NPs name** | **Size by DLS (nm)** | **PdI** | **Size by TEM (nm)** |
|---|---|---|---|
| Bare NPs | 33 ± 2 | 0.13 ± 0.01 | 23 ± 4 |
| 100% coding NPs | 46 ± 1 | 0.11 ± 0.01 | 28 ± 5 |
| 10% coding NPs | 45 ± 2 | 0.12 ± 0.01 | 32 ± 6 |

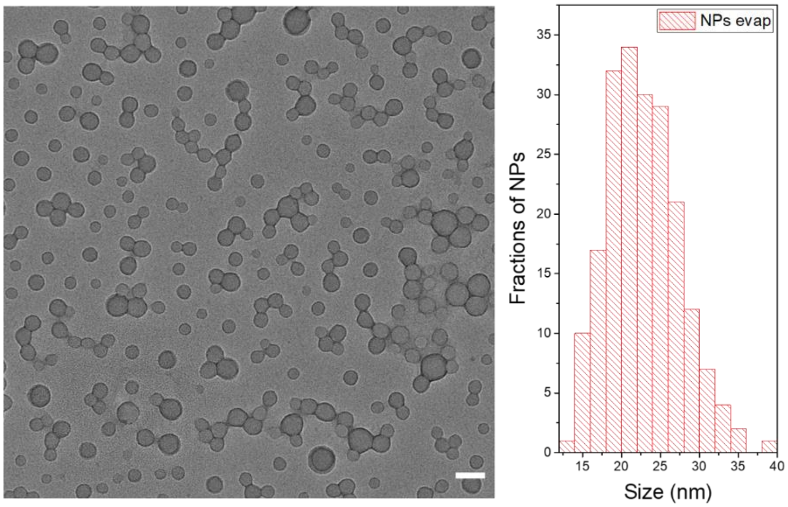


**Figure S1.** TEM image (left) and size distribution diagram (right) of bare PEMA-AspN3 NPs loaded with 33 wt% of R18/F5-TPB dye (with respect to total NP mass). Scale bar: 50 µm.

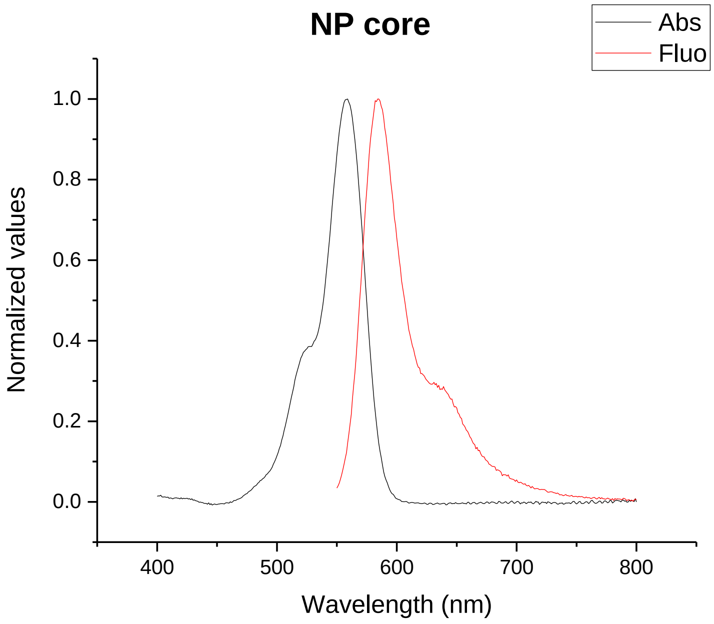


**Figure S2.** Absorption and emission spectra of bare polymeric (PEMA-AspN3) NPs loaded with R18/F5-TPB at 33 wt% (vs total NP mass).

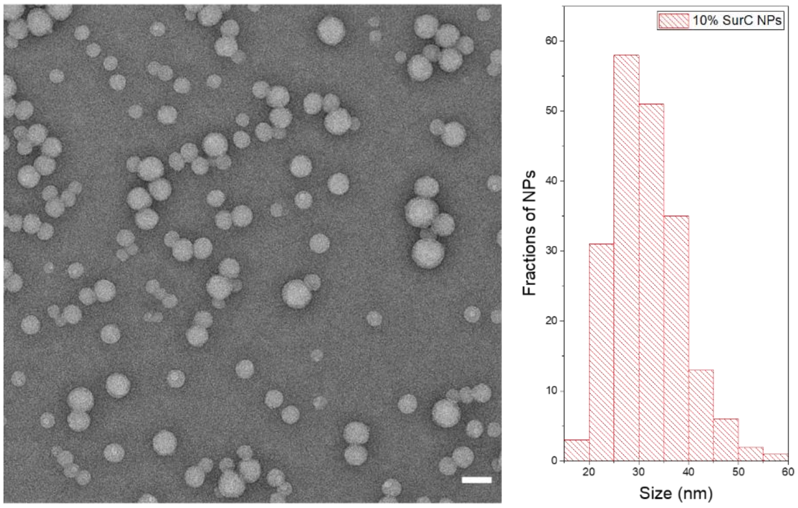


**Figure S3.** TEM image (left) and size distribution diagram (right) PEMA-AspN3 NPs functionalized at 10% with survivin capture coding sequence and 90% A20 (non-coding) sequence and loaded with 33 wt% of R18/F5-TPB dye (with respect to total NP mass). Scale bar: 50 µm.

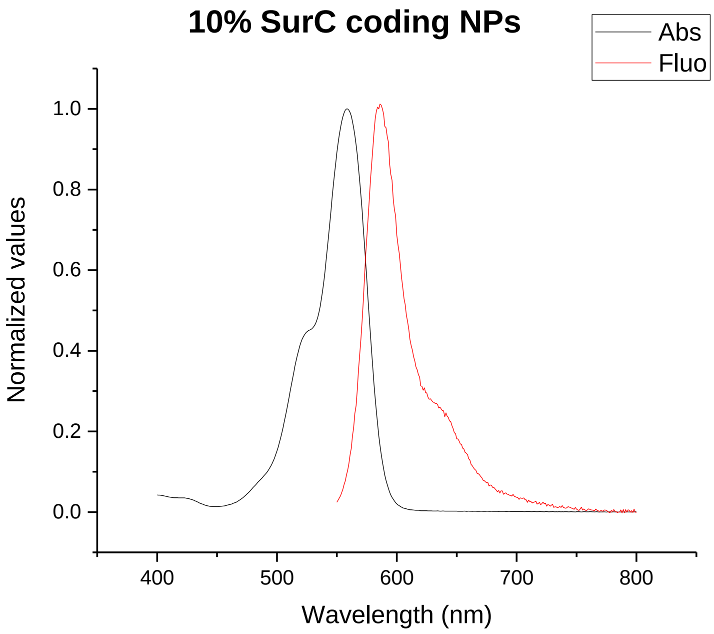


**Figure S4.** Absorption and emission spectra of dye-loaded polymeric NPs coated with 10% SurC and 90% A20 oligonucleotides (10% coding NPs).

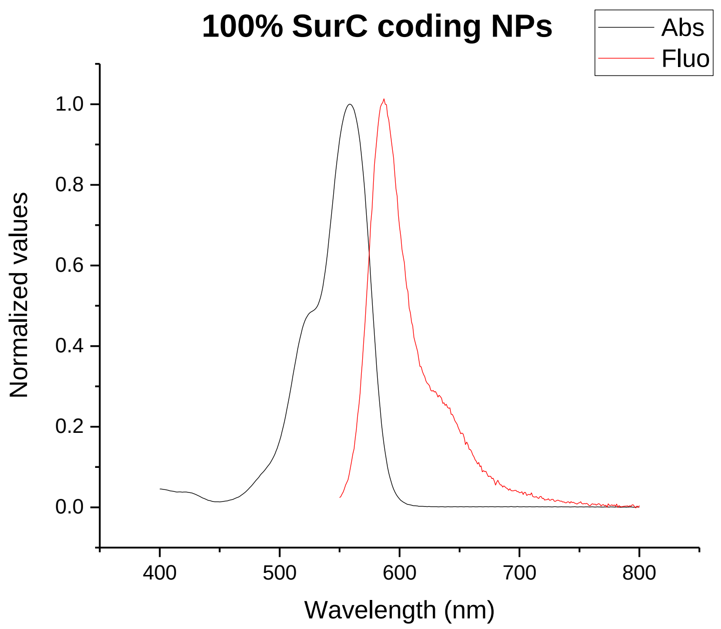


**Figure S5.** Absorption and emission spectra of dye-loaded polymeric NPs coated with SurC oligonucleotide (100% coding NPs).

**THERMAL STABILITY**

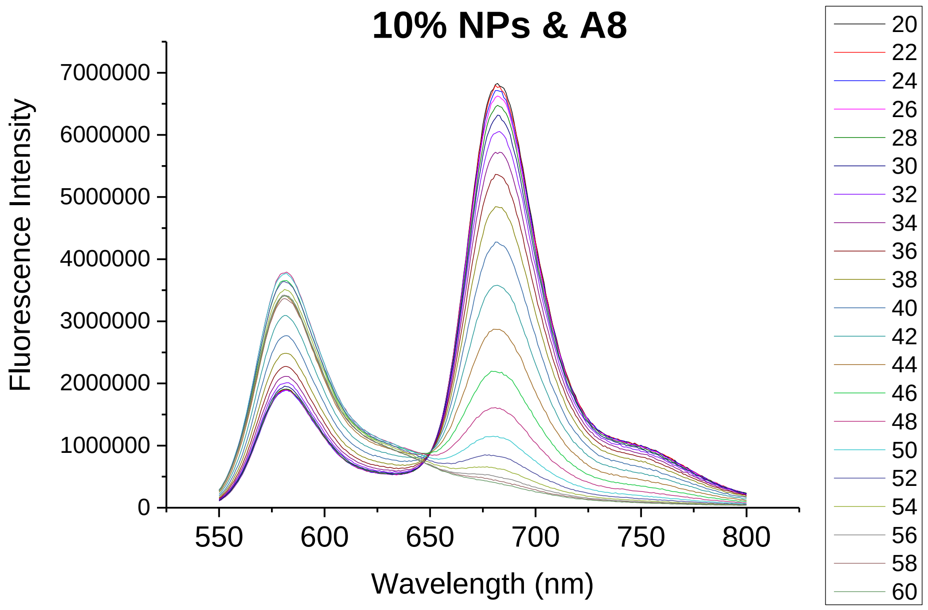


**Figure S6.** Fluorescence spectra of 10% coding NPs mixed with Acceptor-8 at different temperatures.

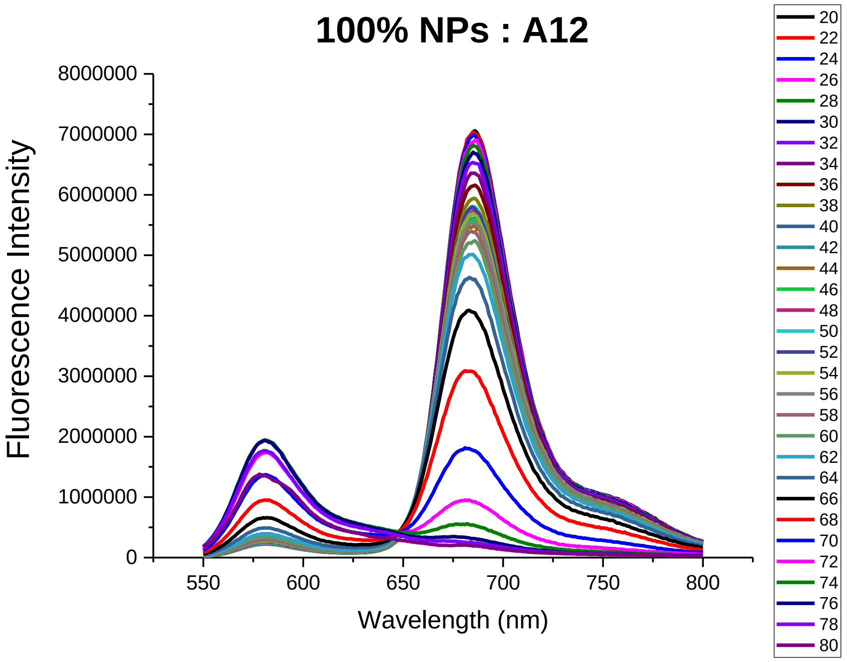


**Figure S7.** Fluorescence spectra of 100% coding NPs mixed with Acceptor-12 at different temperatures.

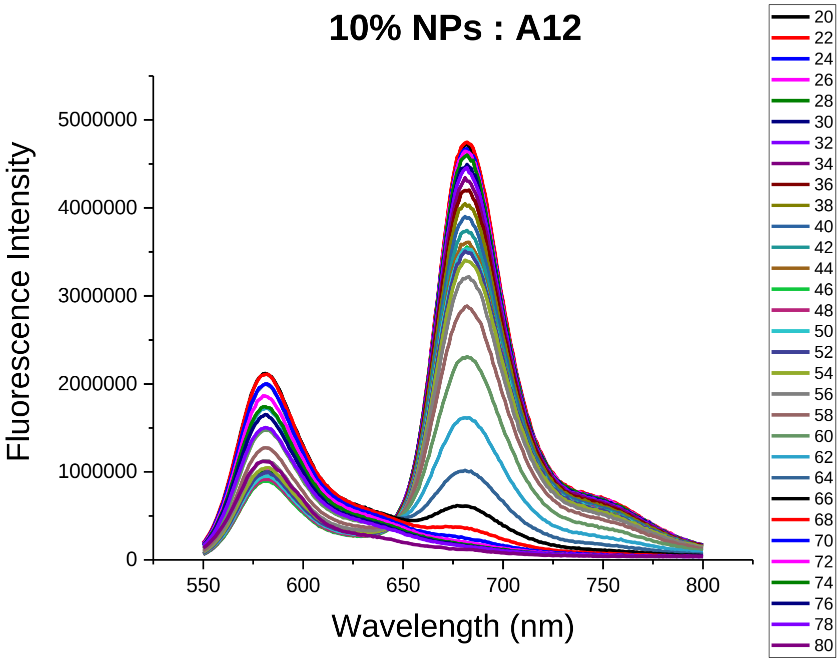


**Figure S8.** Fluorescence spectra of 10% coding NPs mixed with Acceptor-12 at different temperatures.

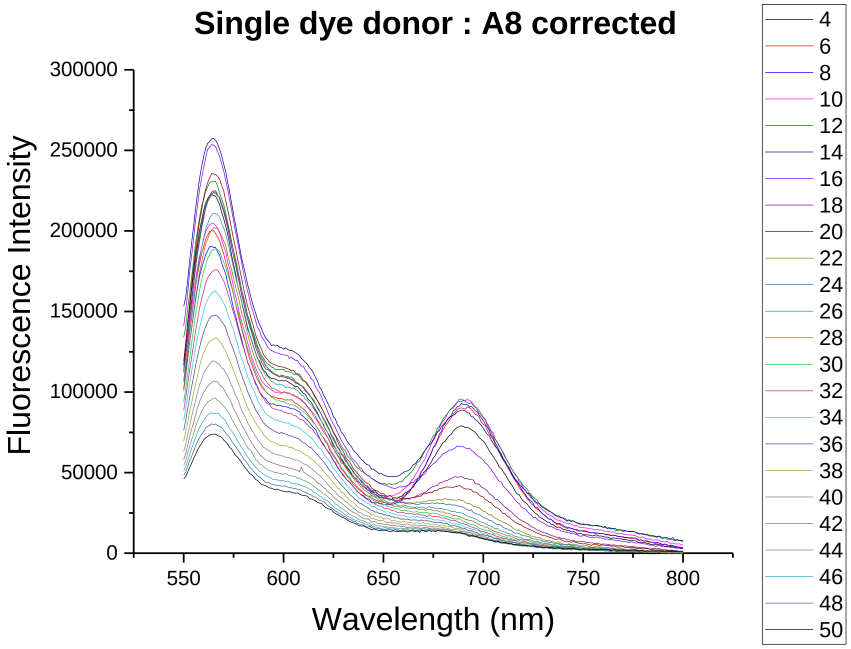


**Figure S9.** Fluorescence spectra of donor dye-labeled SurC oligonucleotide mixed with Acceptor-8 at different temperatures.

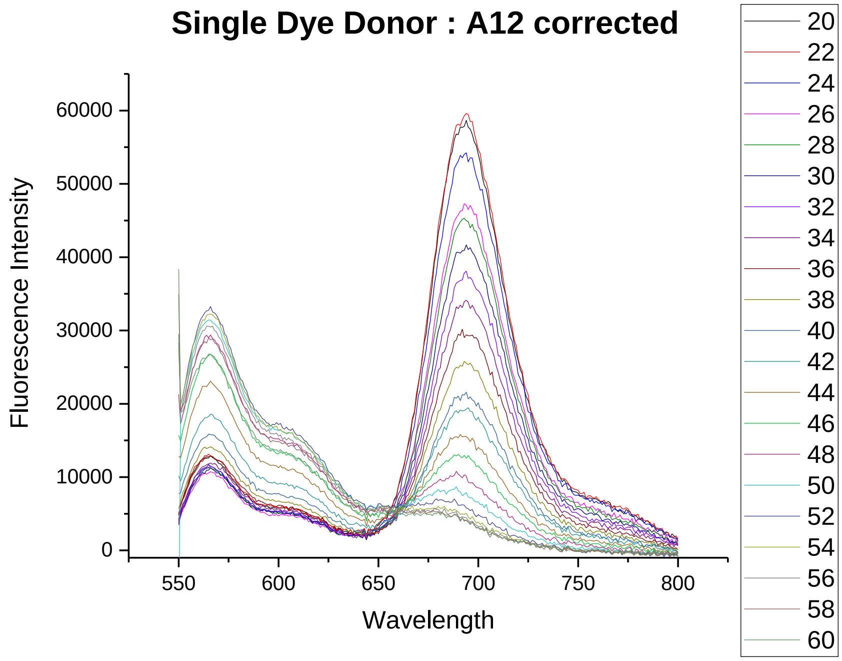


**Figure S10.** Fluorescence spectra of dye-labeled SurC oligonucleotide mixed with Acceptor-12 at different temperatures.

**MUTATIONS**

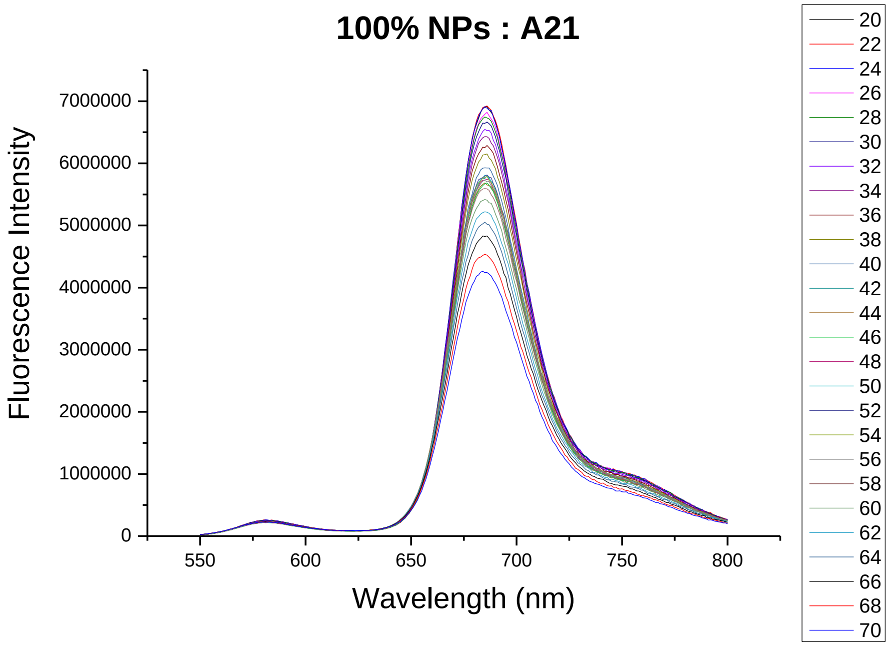


**Figure S11.** Study of mutations with 100% NPs mixed with Acceptor-21: fluorescence spectra at different temperatures.

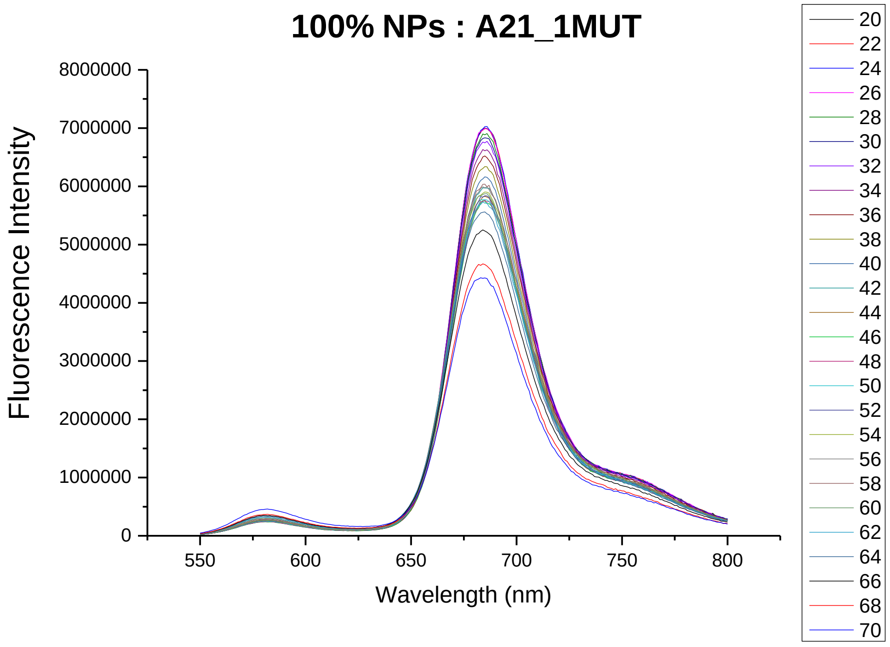


**Figure S12.** Study of mutations with 100% NPs mixed with Acceptor-21 with 1 mutation: fluorescence spectra at different temperatures.

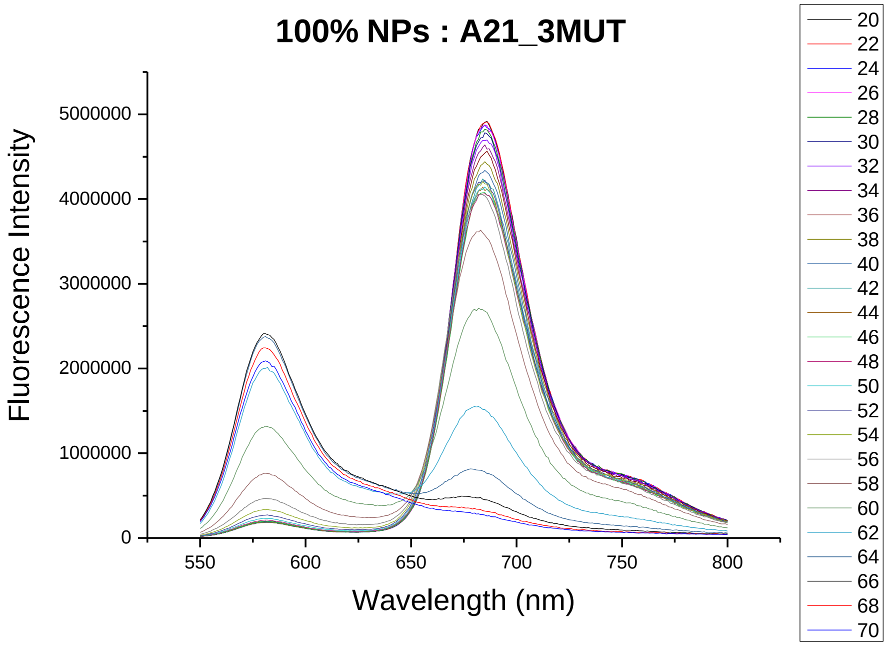


**Figure S13.** Study of mutations with 100% NPs mixed with Acceptor-21 with 3 mutations: fluorescence spectra at different temperatures.

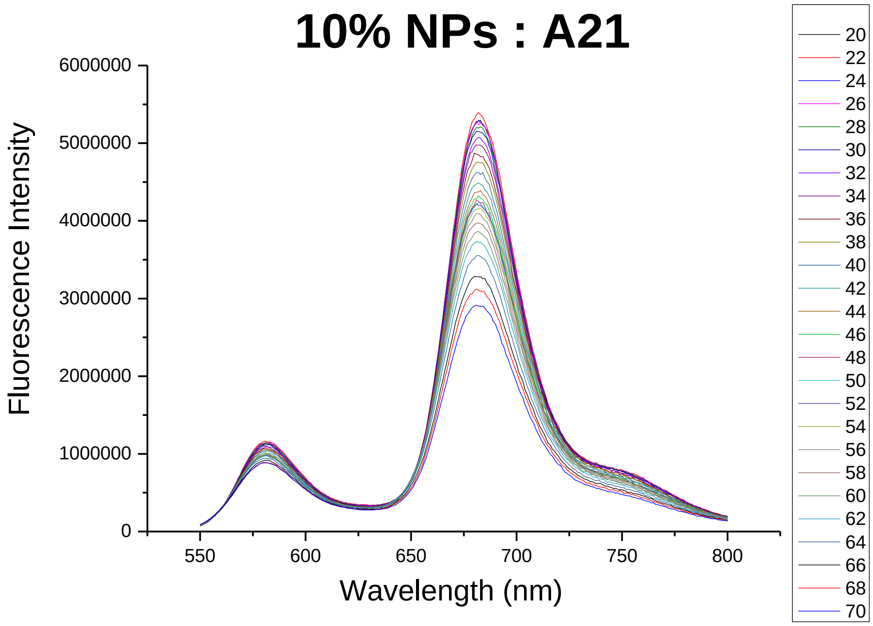


**Figure S14.** Study of mutations with 10% NPs mixed with Acceptor-21: fluorescence spectra at different temperatures.

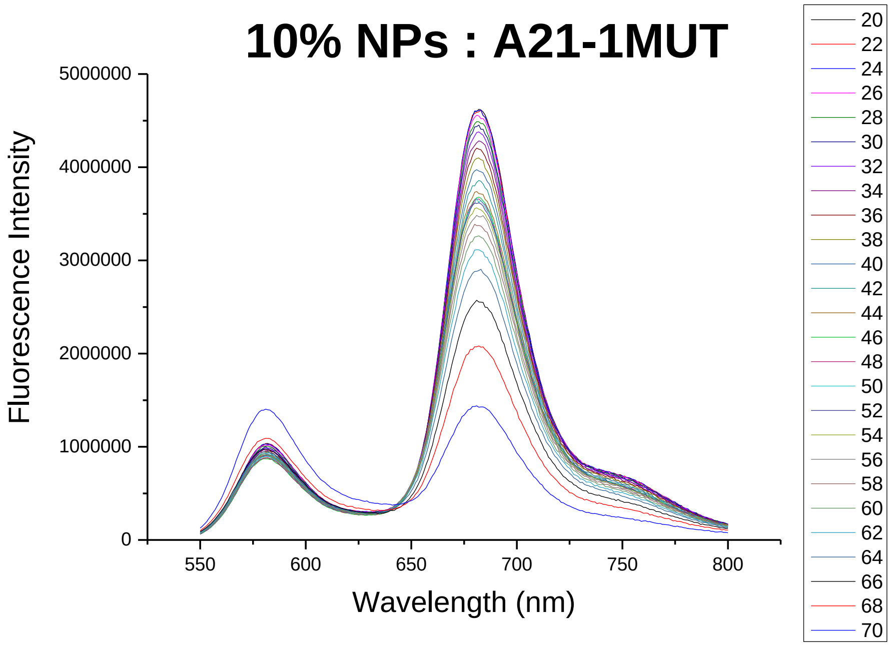


**Figure S15.** Study of mutations with 10% NPs mixed with Acceptor-21 with 1 mutation: fluorescence spectra at different temperatures.

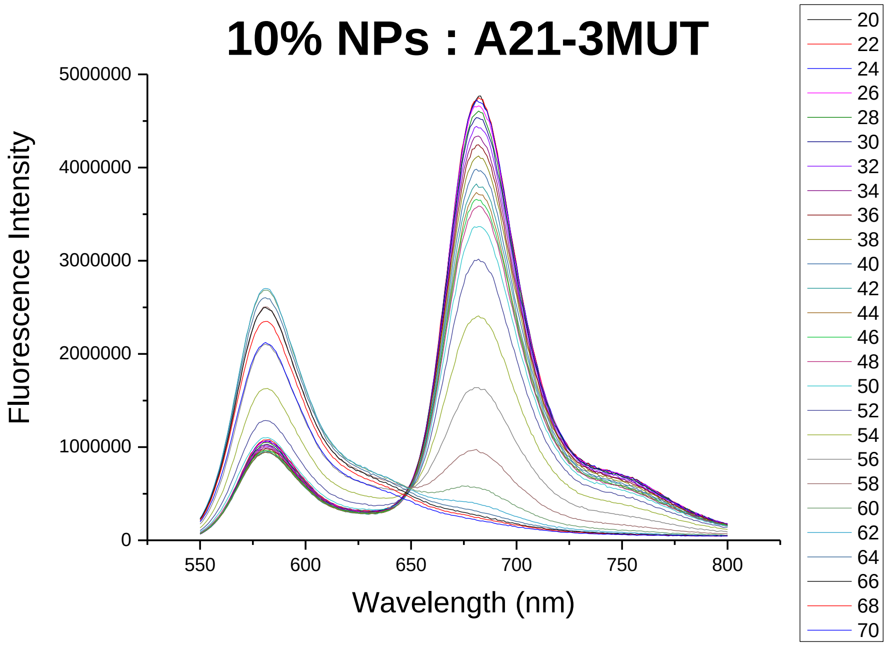


**Figure S16.** Study of mutations with 10% NPs mixed with Acceptor-21 with 3 mutations: fluorescence spectra at different temperatures.

**KINETICS analysis**

**Table S2.** Parameters of the fit for the kinetics curves in Figure S15.

| | | | |
|---|---|---|---|
| Equation | y = Intercept + B1*x^1 + B2*x^2 + B3*x^3 + B4 *x^4 + B5*x^5 | | |
| Weight | No Weighting | | |
| Residual Sum of Squares | 1.05897E-4 | | |
| Adj. R-Square | 0.9912 | | |
| | | Value | Standard Error |
| 1 | Intercept | 0.06656 | 0.00804 |
| | B1 | 0.02618 | 0.0166 |
| | B2 | -0.0027 | 0.0099 |
| | B3 | 5.46212E-4 | 0.00238 |
| | B4 | -6.53053E-5 | 2.46534E-4 |
| | B5 | 2.65807E-6 | 9.2116E-6 |

| | | | |
|---|---|---|---|
| Equation | y = Intercept + B1*x^1 + B2*x^2 + B3*x^3 + B4 *x^4 + B5*x^5 | | |
| Weight | No Weighting | | |
| Residual Sum of Squares | 1.72504E-4 | | |
| Adj. R-Square | 0.99496 | | |
| | | Value | Standard Error |
| 2.5 | Intercept | 0.07625 | 0.01026 |
| | B1 | 0.0762 | 0.02119 |
| | B2 | -0.01544 | 0.01263 |
| | B3 | 0.00205 | 0.00303 |
| | B4 | -1.26987E-4 | 3.14655E-4 |
| | B5 | 2.84518E-6 | 1.17569E-5 |

| | | | |
|---|---|---|---|
| Equation | y = Intercept + B1*x^1 + B2*x^2 + B3*x^3 + B4 *x^4 + B5*x^5 | | |
| Weight | No Weighting | | |
| Residual Sum of Squares | 7.40255E-5 | | |
| Adj. R-Square | 0.99881 | | |
| | | Value | Standard Error |
| 5 | Intercept | 0.10381 | 0.00672 |
| | B1 | 0.08856 | 0.01388 |
| | B2 | -0.01006 | 0.00828 |
| | B3 | 5.28587E-4 | 0.00199 |
| | B4 | 9.69335E-6 | 2.06123E-4 |
| | B5 | -1.34143E-6 | 7.70165E-6 |

| | | | |
|---|---|---|---|
| Equation | y = Intercept + B1*x^1 + B2*x^2 + B3*x^3 + B 4*x^4 + B5*x^5 | | |
| Weight | No Weighting | | |
| Residual Sum of Squares | 1.18032E-4 | | |
| Adj. R-Square | 0.99847 | | |
| | | Value | Standard Error |
| 7.5 | Intercept | 0.14612 | 0.00848 |
| | B1 | 0.09742 | 0.01753 |
| | B2 | -0.0099 | 0.01045 |
| | B3 | 5.86435E-4 | 0.00251 |
| | B4 | -2.1711E-5 | 2.60277E-4 |
| | B5 | 5.27356E-7 | 9.72508E-6 |

| | | | |
|---|---|---|---|
| Equation | y = Intercept + B1*x^1 + B2*x^2 + B3*x^3 + B4 *x^4 + B5*x^5 | | |
| Weight | No Weighting | | |
| Residual Sum of Squares | 5.24884E-5 | | |
| Adj. R-Square | 0.99946 | | |
| | | Value | Standard Error |
| 12.5 | Intercept | 0.13795 | 0.00566 |
| | B1 | 0.1965 | 0.01169 |
| | B2 | -0.04316 | 0.00697 |
| | B3 | 0.00577 | 0.00167 |
| | B4 | -4.05088E-4 | 1.73567E-4 |
| | B5 | 1.13032E-5 | 6.48521E-6 |

| | | | |
|---|---|---|---|
| Equation | y = Intercept + B1*x^1 + B2*x^2 + B3*x^3 + B4 *x^4 + B5*x^5 | | |
| Weight | No Weighting | | |
| Residual Sum of Squares | 1.49949E-5 | | |
| Adj. R-Square | 0.99983 | | |
| | | Value | Standard Error |
| 25 | Intercept | 0.20204 | 0.00302 |
| | B1 | 0.24069 | 0.00625 |
| | B2 | -0.05916 | 0.00372 |
| | B3 | 0.00812 | 8.93953E-4 |
| | B4 | -5.68507E-4 | 9.27697E-5 |
| | B5 | 1.56658E-5 | 3.46628E-6 |

| | | | |
|---|---|---|---|
| Equation | y = Intercept + B1*x^1 + B2*x^2 + B3*x^3 + B 4*x^4 + B5*x^5 | | |
| Weight | No Weighting | | |
| Residual Sum of Squares | 9.36831E-5 | | |
| Adj. R-Square | 0.99876 | | |
| | | Value | Standard Error |
| 37.5 | Intercept | 0.28748 | 0.00756 |
| | B1 | 0.32466 | 0.01562 |
| | B2 | -0.0977 | 0.00931 |
| | B3 | 0.01517 | 0.00223 |
| | B4 | -0.00116 | 2.31881E-4 |
| | B5 | 3.44061E-5 | 8.6641E-6 |
